# A sensorimotor brain circuit for transforming aversive experiences into emotional states

**DOI:** 10.64898/2026.01.15.699823

**Authors:** Li-Feng Yeh, Takaaki Ozawa, Akira Uematsu, Tomoya Duenki, Yuri Ishizu, Andrew J. Murray, Kazuo Okanoya, Joshua P. Johansen

## Abstract

Innately aversive experiences produce emotional states which control behavior as well as long term associative memories^1–3^. Brainstem regions engage innate defensive reactions^1,3–7^, but it is unclear how the nervous system transduces external aversive experiences into higher order, emotional representations in the forebrain to instruct learning and modulate behavior. The prevailing view is that the sensory properties of aversive experiences activate forebrain emotional processing^1,3,8–10^, but classical theories proposed that internal bodily reactions to unpleasant events produce feeling states^11,12^. Here we identify a brainstem cuneiform (CnF) circuit which conveys information about both the external sensory causes and internal motor reactions related to aversive events to the lateral/basal nuclei of the amygdala (LA/B), a brain region which stores emotional memories, to enhance ongoing defensive behaviors and regulate associative memory formation. Glutamatergic CnF neurons project directly to LA/B and receive afferent inputs from aversive-sensory and aversive-motor brain regions. Notably, LA/B neurons encode a sensorimotor state in response to noxious stimuli, with both sensory and motor features transmitted through the CnF-to-LA/B pathway. Finally, optogenetic perturbation experiments showed that during aversive experiences, the CnF-LA/B circuit instructs emotional memory formation and enhances the intensity of ongoing defensive behaviors. These findings reveal how the nervous system constructs an aversive sensorimotor state in the amygdala used for regulating the strength of emotional behaviors and producing memory formation. This demonstrates a clinically relevant brain mechanism for the regulation of emotional processing by both external traumatic events and the accompanying bodily reactions.

## Main

Innately aversive events such as those which are painful or imminently threatening trigger an emotional state comprised of coordinated defensive behavioral, autonomic and hormonal responses^1,3,10,13–15^. In parallel, these experiences activate instructive signaling systems to produce neural plasticity in brain regions which store emotional memories^8,16^, providing the foundation upon which complex emotional representations are built. Brainstem regions are recruited to trigger fast behavioral and visceral responses^1,3,5–7^ and higher order emotion processing centers in the brain such as the LA/B are also engaged to store associative memories^2,9,17,18^ and potentially modulate defensive responses^19,20^. Presently it is unclear what information is encoded in LA/B neurons during innately aversive events or how this information is conveyed there.

Historically, contrasting psychological theories postulated that emotion representations in the brain arise in response to sensory features of innately salient events or that internal bodily responses to these experiences drive emotional responding^11,12,21^. In current animal models of aversive emotion, it is widely assumed that sensory features of innately aversive experiences produce emotional behavioral responding and memory formation^1,3,8–10^. In brainstem structures mediating defensive responses this seems likely, as they receive direct input from sensory systems transmitting aversive information (e.g. nociceptive) and are closely linked to motor control systems in the medulla^1,3^. While the LA/B plays a central role in emotional learning and memory and coordinates behavioral and autonomic defensive response systems^2,9,17,18^, it is not clear what is encoded in these types of higher order emotion processing centers during innately aversive experiences to support learning and other functions.

Supporting the sensory view, LA/B neurons exhibit short latency, sensory-like responses to innate and learned, fear inducing stimuli^22–25^ and activity in LA/B neurons is required for emotional responding to innate and learned aversive events^9,19^. Furthermore, the magnitude of neural plasticity in LA/B neurons occurring during associative learning as well as the strength of subsequent behavioral memory is correlated with the strength of the aversive sensory experience^24,26–28^. However, complicating this sensory based view are findings showing that LA/B neurons also encode motor responses during learned reward and aversive behaviors^29–31^. While the function of this motor coding is unclear, these findings may support earlier theories on the role of internal bodily factors such as defensive behavioral responses to danger (e.g. escape, autonomic responses, etc.)^11,12^ in regulating emotional states.

Resolving this central debate in the emotion field requires an understanding of how the nervous system transforms external innately aversive experiences into emotional state representations in areas like the LA/B to regulate learning and behavior. Here we identify a brainstem circuit which conveys information about both external-sensory and internal-motor features of aversive experience to produce a sensorimotor state in LA/B neurons used for regulating ongoing emotional behaviors and triggering memory formation. This demonstrates a mechanism through which both internal and external aspects of aversive encounters are encoded in forebrain emotional processing centers like the amygdala.

## Amygdala neurons encode both sensory and motor features of innately aversive experiences

We first examined whether noxious stimulation induced responses of LA/B neurons exhibit purely sensory features or encode a sensorimotor state which outlasts the sensory stimulus and reflects both sensory and motor features of the event. Although prior studies reported noxious stimulus evoked responses in LA/B neurons^23,25,28,32^, they did not examine whether LA/B neurons encode purely sensory features of the stimulus or a sensorimotor state which outlasts the sensory stimulus and reflects both sensory and motor aspects of the event. To study this question we quantitatively examined the response properties of LA/B neurons to noxious stimulation using in-vivo electrophysiological tetrode recordings of single LA/B neurons in awake, behaving animals in response to electrical eyelid shocks (**Fig. 1a**, note these recordings were done in parallel with optogenetic manipulations of brainstem CnF inputs to LA/B, see below). Escape behaviors are the dominant response to a variety of innately aversive stimuli including those which are noxious^3,33^. We employed DeepLabCut^34^ to detect and quantify escape responses to shock. We identified LA/B neurons which responded significantly during and/or after shock (**Fig. 1b**) whose activity could correspond to the sensory or motor aspects of noxious stimulation.

**Fig. 1.**
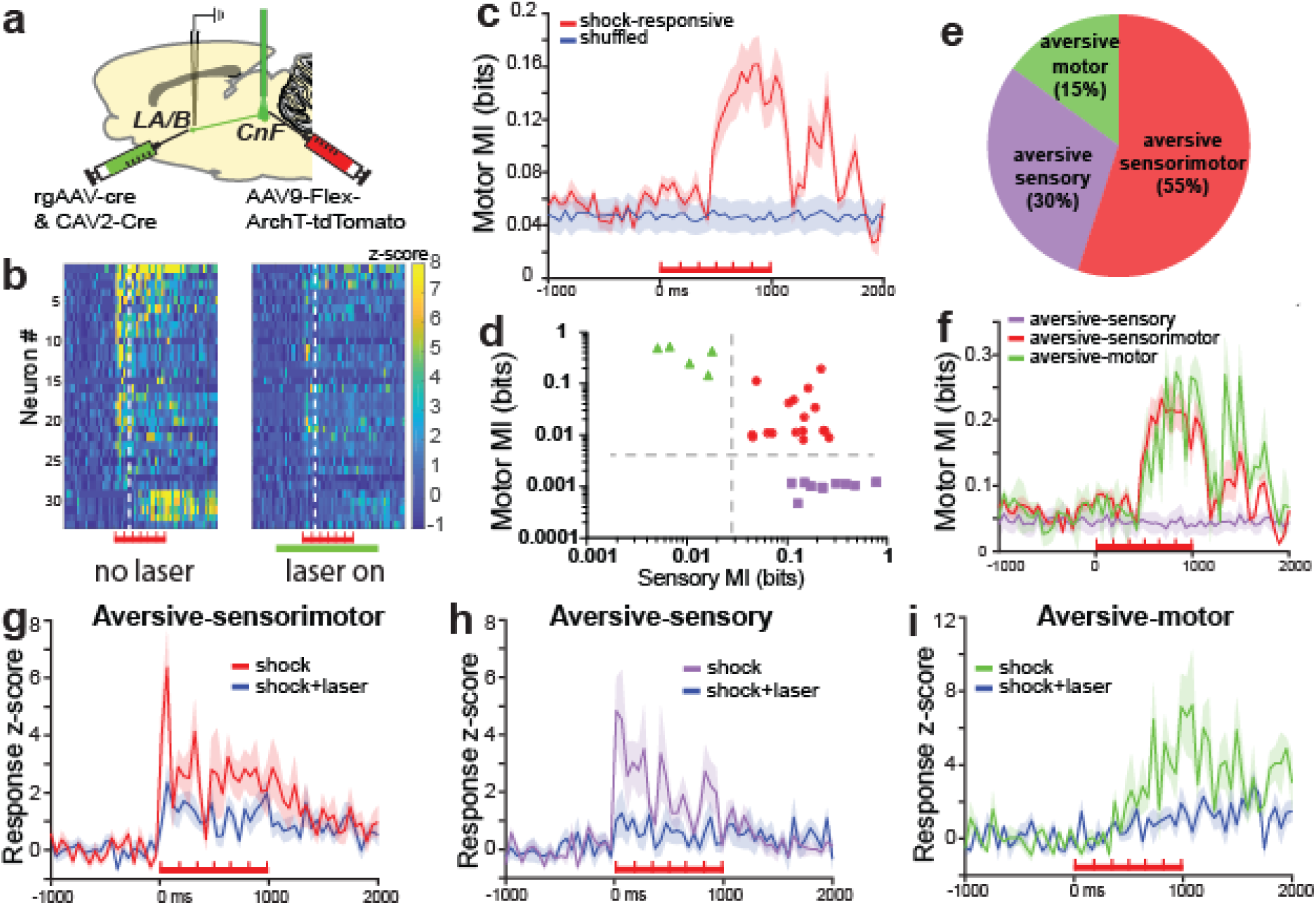
LA/B neurons encode aversive-sensory and aversive motor information which is transmitted from CnF. ***a***) Schematic showing in-vivo single unit electrophysiological recording in LA/B combined with optogenetic inhibition of CnF inputs to LA/B. ***b***) Heat plots showing shock-evoked neural responses in individual shock responsive LA/B neurons (y-axis, n=33/92 cells from 2 animals) without (left) and with (right) laser inhibition of LA/B projecting CnF neurons with time on x-axis (red bars denote 1 s shock periods and pulses, green bar denotes laser on period). White dashed lines indicate average escape onset. ***c***) Peri-event time histogram (PETH) showing the temporal dynamics of mutual information (MI) between LA/B neuronal activity and escape velocity (y-axis, red trace) in response to shock compared with a spike shuffled control (blue trace). x-axis=time in ms. Red bar denotes shock time-period. ***d***) Log plot showing sensory MI (x-axis) and motor MI (y-axis) in single neurons. Colors denote aversive-sensory cells (purple), aversive-motor (green) and aversive-sensorimotor (red) cells. Vertical dotted line=significance threshold for sensory MI while horizontal dotted line=significance threshold for motor MI. ***e***) Pie-chart showing proportions of the different cell types. Motor=15.15% (5/33), Sensorimotor=54.55% (18/33), Sensory=30.3% (10/33). ***f***) PETH showing MI (y-axis) in response to shock in sensory, motor and sensorimotor cell types. x-axis=time in ms. ***g-i***) PETH showing z-score averaged population response of different cell classes without (red, purple and purple traces) or with (blue) CnF inactivation. Inactivation of CnF inputs reduce neural response in all cell types. 1-way ANOVA on spiking activity during 0-2000 ms in laser on and off conditions, F(5, 234) = 19.42, p<0.0001, Bonferroni’s post-hoc analysis comparing laser on vs. laser off periods: Sensorimotor cells: p=0.002, Sensory cells: p=0.0213, Motor cells: p=0.0001). x-axis=time in ms. Data represent mean ± SEM

To disentangle temporally interrelated sensory and motor responses, we utilized a mutual information (MI) approach^35^ which takes advantage of trial-to-trial variability in behavioral and neuronal responses and differences in temporal dynamics within trials. In our paradigm, the escape response onset is delayed relative to shock onset (beginning at 341 +/- 65 ms after shock onset) and the magnitude of shock-evoked behavioral responses vary from trial-to-trial (**Extended Data Fig. 1**). As a result, MI approaches can be used to determine whether the variations in the amplitude of trial-by-trial spiking activity of single LA/B neurons (computed on a fine timescale which captures momentary dynamics or on a broader trial-binned timescale) carries information about the magnitude of the behavioral escape response (compared to a spike shuffled control dataset). We first examined whether the neuronal activity of shock-responsive cells at specific time intervals across the peri-shock period exhibited significant MI with instantaneous escape velocity (‘motor MI’) using a dynamical, sliding MI window (50 ms bins) and relating LA/B spiking activity in single cells and on single trials to the corresponding escape velocity. Averaging MI values for all cells across time, we found that LA/B neuronal activity contained significant MI for escape behavior starting from 470.93 +/- 79 ms following shock onset (**Fig. 1c**). This shows that escape motor information is encoded in the activity of LA/B neurons and reveals the timescale of these dynamics as they relate to behavior. Because the average onset of escape behavior (341 ms, **Extended Data Fig. 1**) precedes the onset of neuronal motor encoding (470 ms, **Fig. 1c**), this suggests that LA/B neural activity reflects, but may not initiate, the aversive motor response.

Next, we used a trial-based MI approach to study single cell encoding. We classified the motor coding properties of each cell by calculating the ‘motor MI’ between their spiking activity and behavioral escape velocity (50 ms bin size). We found that 69.7% of cells significantly encoded motor signals. Next, we calculated ‘sensory MI’ to classify sensory/shock encoding cells, taking advantage of the precise timing of repeatedly presented shock pulses (2 ms pulses, 7 Hz) during the 1 second shock period. Comparing shock on and off periods, we identified LA/B cells as those with significant MI between spiking activity and shock pulse time periods compared with a spike shuffled control (∼85% of peri-shock responsive cells). Three cell types were classified based on their significant sensory MI, motor MI or both (**Fig. 1d-f**). Cells with significant MI for both motor escape velocity and shock were termed ‘aversive-sensorimotor’ (**Fig. 1g**), cells exhibiting only significant sensory MI were labeled ‘aversive-sensory’ (**Fig. 1h**) and cells with significant MI for only motor responses were termed ‘aversive-motor’ (**Fig. 1i**). Aversive motor and aversive-sensorimotor cells exhibited delayed or sustained response dynamics and significant motor MI that outlasted the shock period, while responses in aversive-sensory neurons were limited to the shock period and they did not display an increase in motor MI during the peri-shock period (**Fig. 1f-i**). Together, these results show that LA/B neurons encode both aversive-sensory and aversive-motor information and reveal how sensorimotor state encoding temporally evolves over the course of the aversive experience.

## Identification of a brainstem-amygdala sensorimotor pathway

We next sought to identify a neural pathway to the LA/B which could transmit aversive sensory and/or motor information to generate the sensorimotor state representation in LA/B neurons. While prior studies have identified brain regions which are responsive to noxious or multimodal sensory stimuli and may contribute to aversive processing in LA/B neurons^23,36–39^ and other work has revealed an aversive pathway to the central nucleus of the amygdala^40,41^, a direct instructive input which conveys aversive information to LA/B neurons to regulate learning and behavior has not been identified. Based on our findings that sensorimotor states are encoded in LA/B neurons during innately aversive events, we hypothesized that the aversive signaling pathway to LA/B transmits information about both the sensory aspects of the shock as well as internal information related to escape behavior. Thus, we examined afferent inputs to the LA/B from brainstem sensorimotor regions. Specifically, we studied the CnF because it controls escape responses to aversive stimuli, CnF neurons are both nociceptive and motor responsive and CnF stimulation can modulate forebrain processing^5,7,42–45^. To test whether the CnF projects to LA/B, we injected a retrograde adeno-associated virus (rgAAV)^46^ driving expression of Cre-recombinase (rgAAV-cre) into the LA/B and an AAV expressing a Cre-dependent fluorescent reporter into CnF (AAV-flex-GFP, **Fig. 2a**). In separate experiments, we labeled LA/B projecting CnF neurons using rgAAV2-CAG-H2B-mScarlet immunolabeled CnF neurons with vGlut2, a marker of glutamatergic neurons, and found that these projection neurons were primarily glutamatergic (**Fig. 2b**). Next we examined whether the CnF-LA/B pathway is activated in response to aversive shocks by retrogradely labeling CnF cells which project to LA/B and examining their immediate early gene activity (c-Fos, an index of neuronal activity) following shock presentation. We found that LA/B projecting CnF cells exhibited increased c-Fos activity in the shock treated group relative to a control condition (**Fig. 2c)**. To study whether CnF neurons synapse on LA/B neurons, we next used the enhanced GFP reconstitution across synaptic partners (eGRASP) approach^47,48^ in which pre and postsynaptic component of a split fluorophore were expressed in CnF and LA/B neurons, respectively. Reconstitution of the fluorophore occurs across synapses, allowing us to directly label synaptic connections between CnF and LA/B neurons. We identified clearly labeled puncta, indicative of synaptic connections between CnF axonal inputs and LA/B neurons (**Fig. 2d**). Labeled puncta were preferentially localized to dendritic spines and rarely found on dendritic shafts, further supporting the idea that these are excitatory connections. Descending CnF projections to motor brainstem regions (lateral paragigantocellularis nucleus, LPGi) drive behavioral escape responses^7^. To determine whether the ascending LA/B projecting and descending LPGi projecting pathways are distinct or overlapping, we injected rgAAVs expressing mScarlet (red) or mTurquoise (green) into LA/B and LPGi, respectively (**Fig. 2d**). We quantified the amount of co-labeling in CnF and found largely non-overlapping ascending and descending CnF cell populations. Together, these findings reveal a direct, ascending glutamatergic CnF pathway to the LA/B which is activated by noxious stimuli.

**Fig. 2.**
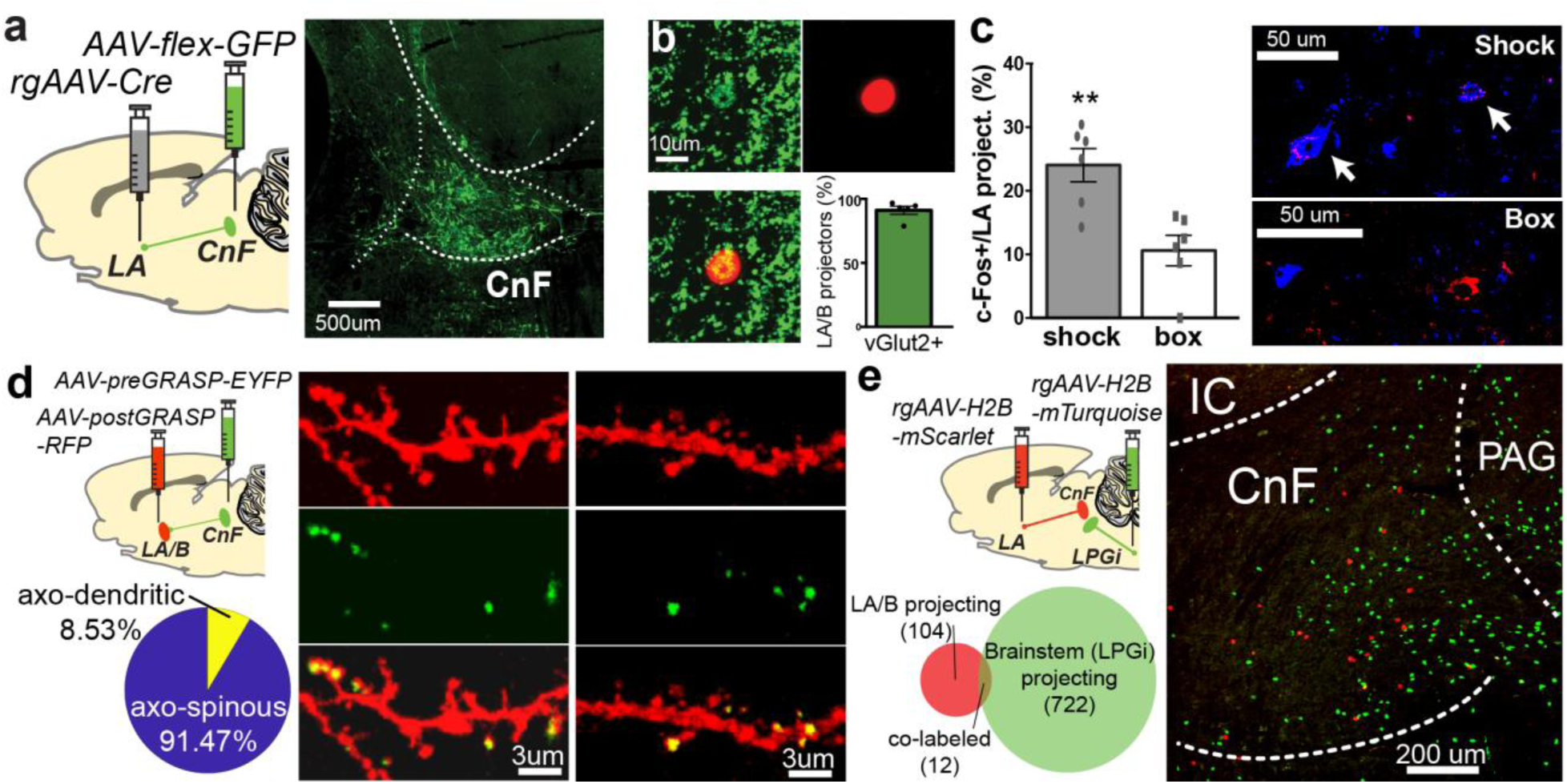
**Anatomical identification of a CnF-LA/B pathway. *a***) Retrograde labeling of CnF neurons projecting to LA/B. Left, schematic showing experimental strategy. Right, labeled LA/B projecting CnF neurons. ***b***) LA/B projecting CnF neurons are glutamatergic. Retrogradely labeled LA/B projecting CnF neurons (red, rgAAV2-CAG-H2B-mScarlet) colabeled with vGlut2+ (green). Quantification of double labeled neurons (bottom right, n=5 rats). ***c***) Footshock (‘shock’) increases c-Fos expression in LA/B projecting CnF neurons compared to animals with ‘box’ exposure alone (quantification in left bar graph, unpaired t-test, t10=3.784, **P = 0.0036). Right example images show retrogradely labeled CnF neurons from LA/B (red), c-Fos positive labeled neurons (blue) in ‘shock’ (top) and ‘box’ (bottom) groups. Arrowheads indicate double labeled cells. ***d***) CnF neurons make synaptic connections with LA/B neurons. Schematic shows eGRASP approach. Pie chart shows proportion of axo-spinous and axo-dendritic synaptic puncta labeling. Example images show red postGrasp labeled LA/B dendrites/spines (top), reconstituted eGRASP labeled synaptic puncta (middle) and overlay (bottom). (n=4 rats, 21 LA/B cells sampled, total axo-spinous puncta = 311, axo-dendritic puncta = 29) ***e***) Dual retrograde labeling of CnF neurons from LA and lateral paragigantocelullar nucleus (LPGi) shows little overlap in these projections. Top left is schematic of procedure. Bottom left shows quantification of the overlap between the two cell populations. Right, representative image showing red (LA projecting) and green (LPGi projecting) CnF neurons. Data represent mean ± SEM.

Using a glycoprotein deleted, transynaptic rabies virus technique^49,50^ (**Fig. 3a**), we next examined the afferent inputs onto the ascending LA/B projecting CnF neurons. Retrograde viruses were injected into the LA/B to express Cre-recombinase in LA/B projecting CnF neurons and cre-dependent helper viruses were injected into the CnF expressing an avian TVA receptor and the rabies virus glycoprotein (AAV9/2-CBA-Flex-TVA950; AAV9/2_AM-CAGGS-FLEX-H2B-HA-P2A-N2c(G)) (**Fig. 3a**). This was followed by injections of a glycoprotein deficient rabies strain (CVS-N2c)^50^ expressing GFP into the CnF. Starter cells expressing both HA-tag (immunolabeled in red) and GFP could be identified in CnF (**Fig. 3b**). We found that CnF-to-LA/B cells receive inputs from a variety of structures (**Fig. 3c-j**) such as midbrain sensorimotor regions and hypothalamic structures important in nociceptive processing, defensive responding and escape behaviors^6,23,28,51–54^ including the periaqueductal gray (PAG) and inferior/superior colliculi as well as the ventromedial hypothalamus. LA/B projecting CnF cells also receive inputs from sensory regions conveying innately aversive information about the external world and visceral functions including the spinal and trigeminal dorsal horn and nucleus tractus solitarius (NTS). Furthermore, ascending LA/B projecting CnF cells receive inputs from sensory and associative cortical areas including the prefrontal, somatosensory and motor cortices as well as basal ganglia regions such as the substantia nigra lateralis (SNl)^55–58^. These anatomical results demonstrate that the ascending CnF-to-LA/B pathway receives afferent inputs from areas involved in sensory, visceral and motor processing related to prepotent, innately aversive events as well as from forebrain decision making and associative cortical structures.

**Fig. 3.**
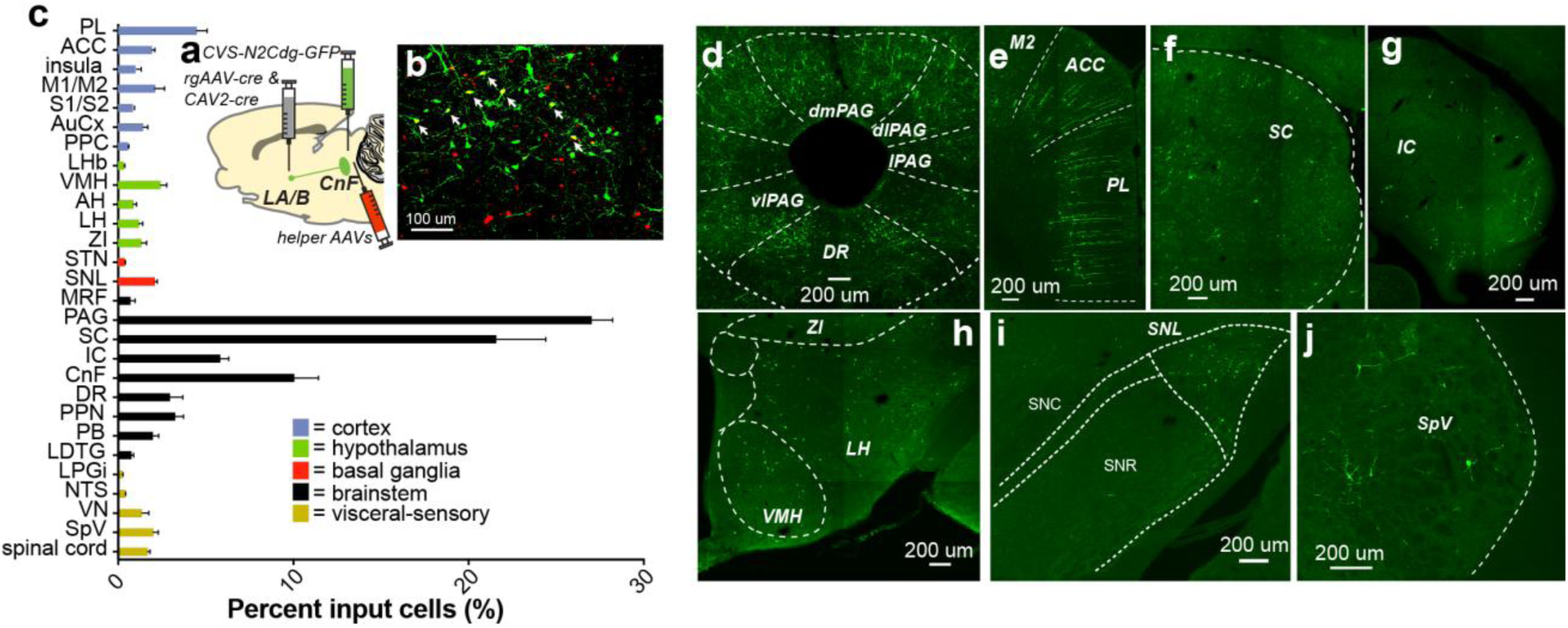
**Whole brain mapping of afferent inputs onto amygdala projecting CnF neurons. *a***) Schematic shows rabies virus transynaptic tracing strategy. ***b***) Image of overlay of helper virus labeled neurons (red), rabies labeled neurons (green) and yellow dual labeled ‘starter cells’ (marked by arrowheads). ***c***) Average individual fractional inputs (relative to all inputs) onto LA/B projecting CnF neurons (n=3). Example images showing labeled input cells from ***d***) Periaqueductal gray (PAG) subregions and dorsal raphe nucleus (DR), ***e***) medial prefrontal cortex including anterior cingulate (ACC), secondary motor (M2) and prelimbic (PL) cortices, ***f***) superior colliculus (SC), ***g***) inferior colliculus (IC), ***h***) ventromedial hypothalamus (VMH), lateral hypothalamus (LH) and zona incerta (ZI), ***i***) substantia nigra lateralis (SNL), compact (SNC and reticulata (SNR), ***j***)spinal trigeminal nucleus (SpV),. Other regions: vestibular nucleus (VN), nucleus tractus solitarius (NTS), lateral paragigantocellularis nucleus (LPGi), laterodorsal tegmental nucleus (LDTG), parabrachial nucleus (PB), pedunculopontine tegemental nucleus (PPN), mesencephalic reticular formation (MRF), subthalamic nucleus (STN), anterior hypothalamus (AH), lateral habenula (LHb), posterior parietal cortex (PPC), auditory cortex (AuCx), primary/secondary somatosensory cortex (S1/S2). Data represent mean ± SEM.

## The cuneiform nucleus conveys aversive sensorimotor information to amygdala

Having identified an anatomical CnF-to-LA/B pathway which is activated during or after noxious stimulation and receives input from aversive sensory and motor systems, we next tested whether this circuit conveys aversive-sensory, aversive-motor or both types of information to the LA/B. We did this by optogenetically inactivating LA/B projecting CnF cells during the peri-shock period and quantifying the effect of the inactivation on aversive-sensory and aversive-motor responding in the same LA/B neurons recorded above (**Fig. 1**). We injected a cocktail of two retrograde viruses expressing Cre-recombinase into the LA/B (rgAAV-Cre/canine adenovirus-cre (CAV2-Cre)^59^) cocktail) and a virus encoding a Cre-dependent inhibitory opsin Archaerhodopsin-3.0T (AAV-CAG-flex-ArchT-tdTomato)^60,61^ into the CnF. We then optically inhibited LA/B projecting CnF neurons during the peri-shock period and determined the effect of this manipulation on LA/B neural coding. Examining all LA/B neurons activated during the peri-shock period, we found that shock-evoked responding was significantly reduced (**Fig. 1b**). We then studied the effects of CnF-LA/B circuit inactivation on the responses of the MI identified subpopulations of cells. We found that inactivation of this pathway reduced responding in aversive-sensory, aversive-motor and aversive-sensorimotor cell populations and attenuated both sensory and motor components of the shock evoked response (**Fig. 1f-h**). This shows that CnF conveys both aversive-sensory and aversive-motor information to the LA/B underlying the sensorimotor state representation in LA/B neurons.

## Cuneiform-amygdala pathway regulates aversive memory formation

Our results thus far reveal a CnF-to-LA/B circuit for generating an aversive sensorimotor representation in LA/B neurons. We next examined the behavioral function of this pathway and the sensorimotor representation it supports. Specifically, we examined whether it regulates aversive memory formation. To study this question, we used an associative learning paradigm termed auditory fear conditioning (**Fig. 4a**) in which animals learn that a neutral auditory stimulus predicts the occurrence of a noxious stimulus (electrical shock). In response to the auditory stimulus following learning, animals exhibit behavioral and visceral responses, including freezing behaviors (the dominant learned behavioral response), which reflect implicit aspects of aversive emotional memory^62^. Given the importance of LA/B in aversive associative learning and the fact that LA/B neurons encode both sensory and motor aspects of innately aversive events, we used this behavioral paradigm to first ask whether the CnF-LA/B pathway is necessary for aversive learning. To do this, we first retrogradely infected CnF-to-LA/B projecting neurons using a viral approach (**Fig. 4a**). A retrograde rabies virus variant^63^ which is taken up directly by synaptic terminals was injected into LA/B to infect LA/B projecting CnF neurons with the inhibitory archaerhodopsin-T3.0^61^ or eGFP as a control (RVG-eArchT3.0/eGFP) and optical fibers were implanted above the CnF to inhibit these cells during the peri-shock period of fear conditioning (‘ArchT-overlap’ group). The effect of the inactivation in this experimental group was compared to two control groups (‘eGFP’ group and ‘ArchT-offset’ where CnF-LA cells were inhibited during the inter-trial interval instead of during the peri-shock period). Inactivation of the CnF-LA/B pathway neurons abolished aversive memory formation measured 24 hours after learning (**Fig. 4b**). Second, to determine whether shock-evoked activity in synaptic inputs to LA/B from CnF is necessary for fear learning, we retrogradely infected CnF-LA/B neurons with Cre-recombinase using a combination of CAV2-Cre and rgAAV-Cre followed by a Cre-dependent archaerhodopsin or tdTomato AAV (AAV9-CAG-Flex-ArchT-Tdtomato, AAV9-CAG-flex-tdTomato) into the CnF which produced robust fiber labeling in LA/B (**Fig. 4c**). We then implanted optical fibers above LA/B to inhibit neurotransmitter release from CnF-LA/B neurons and repeated the same experiment described above. We found that inhibition of this pathway reduced aversive memory formation (**Fig. 4d**). By contrast, inactivation of the CnF terminals in LA/B during memory recall had no effect on behavioral freezing responses (**Extended Data Fig. 2a-c**). Together this shows that activity during the peri-shock period in CnF-LA/B projecting cells or their nerve terminals in LA/B is necessary for forming aversive memories, but not for expressing conditional freezing behavior after learning has occurred.

**Fig. 4.**
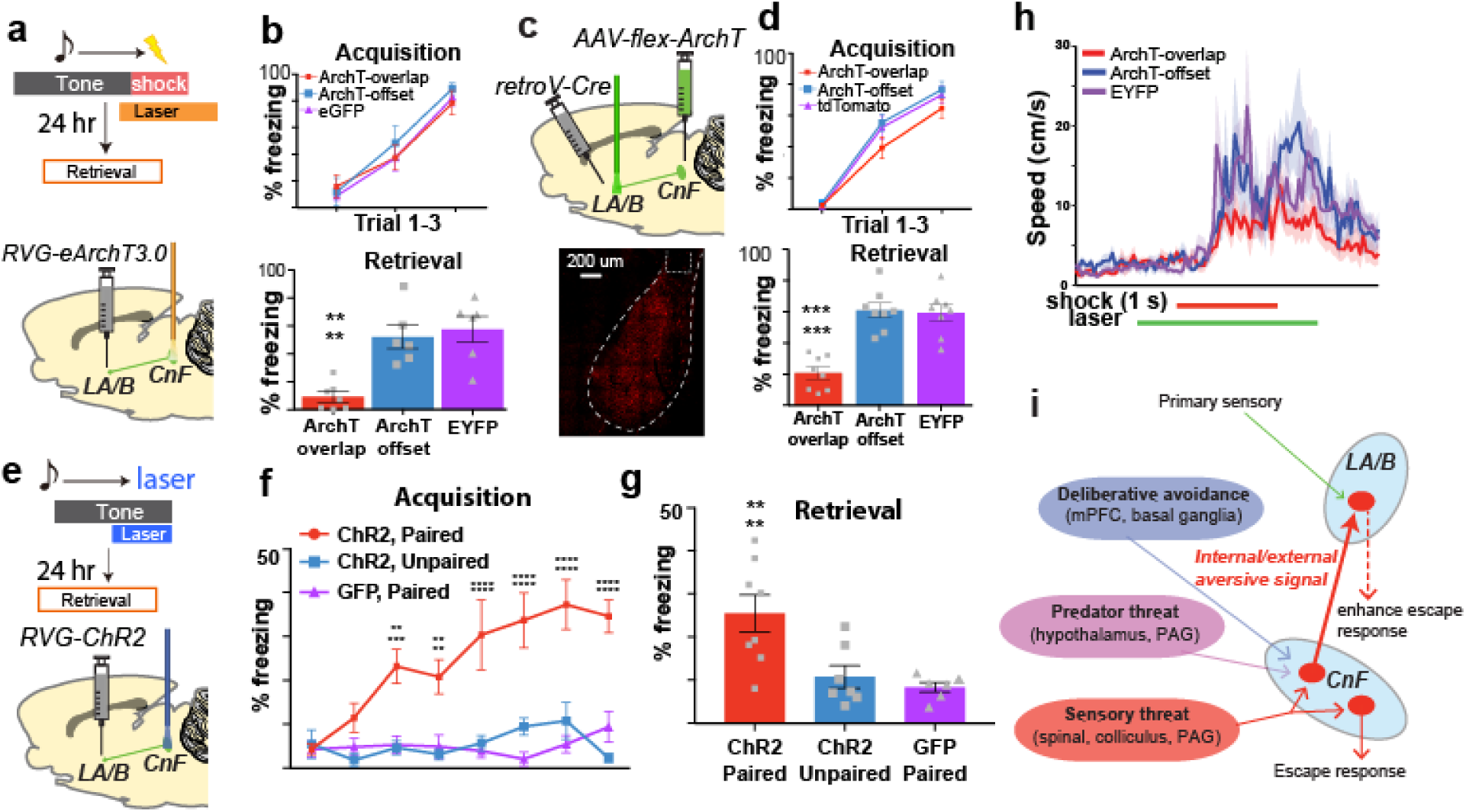
**CnF-LA/B pathway instructs aversive memory formation. *a***) Fear conditioning paradigm with laser inhibition during the peri-shock period (top) and viral/optogenetic strategy (bottom). ***b***) Optogenetic inactivation of CnF-LA/B projecting cell bodies reduces auditory fear memory formation. Top, acquisition. Bottom, memory retrieval (one-way ANOVA, F (2, 16) = 14.61, *P*=0.0002). ***c***) Optogenetic inhibition of CnF terminals in LA/B viral/optogenetic approach (top) and fluorophore labeled CnF terminals in LA/B (bottom). ***d***) Optogenetic inactivation of CnF terminals in LA/B reduces auditory fear memory formation. Top, acquisition. Bottom, memory retrieval (one-way ANOVA, F (2, 21) = 24.34, *P*<0.0001). ***e***) Conditioning paradigm for tone-laser stimulation of Cnf-LA/B pathway (top) viral/optogenetic strategy (bottom). ***f***) Pairing tone with optogenetic stimulation of CnF-LA/B pathway produces fear learning during the acquisition session (two-way ANOVA, F (2, 144) = 76.64, *P*<0.0001). Single trials on y-axis. ***g***) Increased conditioned freezing is maintained in ChR2 paired group at memory retrieval (one-way ANOVA, F (2, 18) = 8.471, *P*=0.0026). ***h***) Effect of optogenetic inhibition of CnF-to-LA/B pathway on escape behaviors. ***i***) Interpretative working model showing CnF-to-LA pathway receiving inputs from brain regions involved in aversive sensory processing and hierarchically organized defensive responding and, in turn, sending external-sensory and internal-motor information to LA/B to instruct associative emotional memory formation and enhance escape responding. \*\**P*<0.01, \*\*\**P*<0.001. Data represent mean ± SEM

To determine whether activity in this pathway is sufficient to produce aversive learning, we expressed the excitatory channelrhodopsin^64^ (ChR2) in LA/B projecting CnF cells and paired an auditory stimulus with direct optical stimulation of this pathway in the absence of shock (**Fig 4e**). During learning, animals receiving tone-optical stimulation pairings (‘ChR2-paired’) began to freeze to the auditory tone over the course of training, but this did not occur in animals expressing GFP (“GFP-paired’) or in animals in which the tone and optical stimulation were unpaired (‘ChR2-unpaired’) (**Fig. 4f**). This learning was maintained 24 hours later (**Fig. 4g**), demonstrating that tone-stimulation pairings produced a long-term memory. Together, these results support the hypothesis that the CnF-LA/B pathway is necessary and sufficient to produce aversive learning.

## Cuneiform-amygdala pathway modulates ongoing escape behavior

Our results show that the CnF-LA/B pathway functions to support aversive learning, but it is unclear whether this pathway also regulates defensive escape behaviors during innately aversive experiences. Given the importance of the descending CnF-medulla pathway in producing fast, escape-like motor responses^7^, we next examined whether the ascending CnF-LA/B pathway participates in escape responding. We found that stimulating the CnF-LA/B pathway did not produce escape or increases in locomotor activity, but did engage flat-back approach and rearing responses (**Extended Data Fig. 3d-g**), behaviors which have been classified as defensive responses in ethological studies^65^. We next examined the effect of inhibiting the terminals of CnF cells in LA/B on escape responses to shock. We found that animals initiated escape responding normally, but the magnitude of these responses was blunted (**Fig. 4h, Extended Data Fig. 3h-j**). Together these data show that this ascending CnF pathway does not drive escape behaviors directly, but boosts ongoing escape through projections to the LA/B.

## Discussion

Innately aversive experiences such as those which are painful or threatening (e.g. noxious stimulation, looming stimuli, etc.) activate neural circuits to engage a concerted emotional response involving behavioral, autonomic and hormonal sequelae as well as neural instructive signaling pathways which trigger memory formation^1,3,8,10,13–16^. While brainstem systems such as the CnF are thought to directly produce defensive behavioral responses^1,3,5–7^, how aversive information is conveyed to and used by forebrain structures involved in higher order emotional representations and learning has been unclear. Here, we identified a brainstem pathway which provides information about the external sensory properties and internal bodily reactions to innately aversive experiences to construct a sensorimotor representation in the LA/B (**Fig. 4i**). This pathway functions to produce aversive memory formation but, unlike descending motor pathways from the CnF^5,7^, activity in the ascending CnF-amygdala pathway does not recruit escape responses directly. Instead, it conveys information about ongoing escape behaviors to the amygdala and is necessary to enhance the amplitude of longer lasting behaviors. These findings suggest a modification of existing psychological theories of emotion by demonstrating that higher order, aversive representations arise in response to both the external sensory features and the internal bodily reactions to innately aversive experiences. Furthermore, these results identify a biological substrate for this process involving the CnF, an early brainstem sensorimotor region, which conveys information about the sensory causes and behavioral reactions to noxious stimulation to forebrain emotional processing areas in the amygdala.

The behavioral/visceral responses to innately aversive events as well as the neuronal representations underlying these responses are temporally evolving and dynamic. The initial component of the behavioral response to innately aversive stimuli is escape, and we show here that the neural representation during these encounters is encoded as a sensorimotor state in the amygdala driven by bottom-up influences from the brainstem. However, emotional states are longer lasting and change over time. Furthermore, defensive responses are organized according to the imminence of threat, learning and cognitive evaluations^1,10,66,67^. For example, after the initial flight response to noxious stimuli, freezing behaviors emerge. This behavior is not an unconditioned response as it require some form of working memory or associative learning between the aversive stimulus and contextual features of the environment^33,68^. Thus, the longer lasting features of emotional state, which incorporate contextual features and prior, learned information, may be mediated by top-down influences from areas like the amygdala to the brainstem. Other types of aversive experiences such as those which are learned or internally generated may be triggered by sensory stimuli in a top-down way, engaging forebrain systems first which then produce defensive responding through projections to the brainstem^9,69–72^. In this situation, information about motor and autonomic activity could then be conveyed back to forebrain regions to modulate ongoing representations there. Consistent with this idea, previous studies found that learned freezing responses are augmented by encoding of respiration in prefrontal circuits^73^, that parabrachial projections convey breathing information to CeA to regulate anxiety^74^ and that insular cortex neurons can be modulated by interoceptive cues during extinction of fear responding^75,76^. Our results provide a neural circuit-based framework for future studies of emotional processing involving hierarchical and dynamic reciprocal interactions between brainstem and forebrain structures for initiating and fine-tuning temporally evolving innate, learned and internally generated emotional states.

Many brainstem regions are sensorimotor structures, receiving primary sensory input and sending axonal projections to spinal skeletomotor and autonomic control systems^3,4,54,77^. These brainstem sensorimotor regions also send ascending projections to forebrain regions like the amygdala^2,45,54^, but the role of these ascending projections as they relate to sensorimotor function is not clear. Our findings suggest a novel framework for understanding the role of brainstem systems in controlling behavior and modulating forebrain functions such as emotional processing and learning. Specifically, these results suggest that sensorimotor brainstem areas can integrate exteroceptive sensory information related to salient experiences and send segregated ascending and descending projections for controlling forebrain and motor processes, respectively. Our anatomical evidence indicates that the ascending CnF pathway receives inputs from regions which participate in distinct, hierarchically organized defensive responses. Specifically, ascending CnF neurons are innervated by brain areas involved in escape behaviors evoked by innately aversive nociceptive, looming and startling auditory sensory stimuli (spinal cord, PAG, colliculus) as well as responses to predator and social threats (hypothalamus, PAG)^6,23,28,51–54^. The CnF-LA/B projecting cells also receive synaptic inputs from forebrain regions (prefrontal cortex, motor cortex, basal ganglia) which contribute to motor planning and more deliberative and learned defensive processing such as active avoidance^55–58^. This information could set emotional state coding, instruct aversive learning and enhance ongoing emotional responses through ascending CnF projections to forebrain regions while descending CnF-medullary projections generate motor escape or avoidance responses. The fact that descending motor production and ascending emotional regulating CnF projections are segregated suggests that these functions can be differentially controlled.

These findings also have important implications for understanding trauma related psychiatric conditions. Only ∼20% of individuals exposed to traumatic experiences go on to develop post-traumatic stress disorder (PTSD)^78,79^. Defining symptoms of PTSD include exaggerated fear learning and memory, difficulty in extinguishing traumatic memories and heightened behavioral reactivity to aversive, startling or even innocuous stimuli^78–81^. One possibility raised by the present results is that the magnitude of internal-bodily reactions to directly experienced or recalled traumatic experiences underlie individual differences in the propensity to develop and maintain PTSD symptoms. Understanding the neural mechanisms through which bodily factors influence aversive learning, memory and behavior may provide new insights into the etiology and persistence of PTSD and point the way toward novel treatments for this disorder.

## Methods

### Subjects

Male Long Evans rats weighing 250-375g were singly housed on a 12 hr light/dark cycle (lights on at 7 a.m. to 7 p.m.) in the vivarium which was maintained at a constant temperature (23 °C) and given food and water ad libitum. Experimental procedures were approved by the Animal Care and Use Committees of the RIKEN Center for Brain Science.

### Stereotaxic surgeries

Standard surgical protocols were performed for all stereotaxic injections/implantations. Briefly, rats were anesthetized with isoflurane (3-5% for induction and 1.5%-3% for maintenance) and placed in a stereotaxic frame (Kopf Instruments). The injection /implantation sites were located based on distance from bregma on the skull. Viruses or tracers were delivered with stainless steel injection cannulae (26 gauge, Plastics One), which were connected to 1 mL Hamilton syringe with polyethylene tubing. After virus-loaded cannulae were advanced to the target location, a syringe pump (PHD2000, Harvard Apparatus) was used to continuously infuse virus at 0.07μl/min flow rate. The cannula was maintained at the injection site for 15 minutes after the infusion was completed, then slowly withdrawn. For the in-vivo optogenetic experiments, the optical fibers held in ferrules (200 μm core, N.A. = 0.37, Thorlabs) were implanted in the same surgery. The ferrules were affixed to the skull with acrylic dental cement. For the anatomical tracing experiments, the surgical incision was sutured after virus/tracer injection. For the in-vivo electrophysiology experiments, incisions were sutured after virus injection. Then, following a 6 week incubation period, a separate surgery was performed to implant the optical fibers and recording electrodes.

### Immediate-shock induced c-Fos expression

Rretrograde tracer (red retrobeads, Lumafluor Inc.) was injected into LA/B (AP: -3.1mm, ML: ±5.4mm, DV: -8.6mm for 0.3μl). One week after surgery, animals were randomly assigned to ‘shock’ or ‘box’ group. The behavioral session was 5 mins long. The ‘shock’ group animals received 3 scrambled footshocks (1s, 0.7mA, inter-stimulus interval: 1s) immediately after the animals were put into the behavioral boxes. The ‘box’ group animals were placed in the behavioral boxes but did not receive footshocks. Each animal was then returned to the animal housing area after the behavioral session and sacrificed 90mins later. The tissue sections throughout the anterior-posterior extent of CnF (AP: -7.4 mm to -9 mm) were collected and the immunohistochemistry procedures were performed on sections 160 μm apart. The number of c-Fos immunoreactive cells which were co-localized with retrobeads were counted using a confocal microscope (Olympus FV-3000).

### In vivo optogenetic manipulations during behavioral experiments

For the optogenetic cell-body inactivation of LA/B projecting CnF neurons, the RVΔG-eArchT3.0-EGFP or RVΔG-EGFP was bilaterally injected into LA/B (AP: -3.0mm, ML: ±5.3mm, DV: -8.6mm, 0.4μL total injection volume) and the optical fibers were bilaterally implanted above CnF (AP: -8.4mm, ML: ±3.85mm, DV: - 4.8mm with 20° angle). For the optogenetic cell-body stimulation experiment, the RVΔG-ChR2-EGFP or RVΔG-EGFP were injected into LA/B and fibers were implanted above CnF, at the same location as in the cell-body inactivation experiments. Behavioral experiments began 4 days after rabies transfection. For the optogenetic terminal inhibition experiment, the CAV2-cre and rgAAV2-mCre (mixed in 1:1 ratio by volume) were bilaterally injected into LA/B (AP: -3.0mm, ML: ±5.3mm, DV: -8.6mm for 0.4μl); and AAV9-CAG-FLEX-ArchT-tdTomato or AAV9-CAG-FLEX-tdTomato was bilaterally injected into CnF (AP: -8.4mm, ML:±3.85mm, DV: -5.3mm with 20° angle, 0.3 μL total injection volume). Optical fibers were implanted bilaterally in LA/B (AP: -3.0mm, ML:±5.3mm, DV: -7.2mm). The animals underwent behavioral experiments after an 8 week incubation period following virus transfection. In the optogenetic inhibition of LA/B-projecting CnF neurons experiments (Fig. 4a), orange laser light (589nm, Shanghai Laser) was delivered continuously during peri-shock periods (from 400ms before shock onset to the 400ms after US termination). In the terminal inhibition experiments (Fig.4d), green laser light (532 nm, Laserglow Technologies) was delivered with the same onset/offset timing as described above. In the stimulation experiment, the blue laser (473nm, Shanghai Laser) pulses (5ms, 20Hz) were delivered for 10 seconds, beginning 10 sec after auditory CS onset and co-terminating with the CS. For all optogenetic studies, we verified that the laser intensity was 15–20 mW from the tips of optical fibers before each experiment using a laser power meter.

### Fear conditioning

Animals were randomly assigned to experimental groups before each experiment. They received 2 days handling by experimenters ahead of the conditioning sessions. For all auditory fear conditioning in behavioral studies, animals were placed into a sound-isolating chamber (Med Associates) and received auditory CSs (74 dB, 5-kHz tone pips at 1 Hz with 250 ms on and 750 ms off for 20 s) and electric shock unconditioned stimuli (US) (1 sec, 0.7mA) which co-terminated with the CS. All trials were separated with variable inter-trial intervals (2.5 min on average). For optogenetic inactivation during fear conditioning experiments, animals were conditioned with three CS–US pairings, and procedures were identical for all groups. Trained animals underwent a retrieval test 24hrs after conditioning. During retrieval, animals received five CS-alone presentations in a different context. For optogenetic inactivation during fear retrieval, animals were re-conditioned with two CS-US pairings in the original conditioning context. 24 hr after re-conditioning, animals received 4 CS-alone presentations in the retrieval context. Two of the retrieval CSs were presented with green laser (laser onset 400 ms before CS onset, laser offset 400 ms after CS offset), while the other two were presented without laser. The order of ‘Laser-on’ and ‘Laser-off’ trials were counterbalanced across animals (i.e. in half of animals, CS trial 1 &2 had laser while in other animals laser occurred during trials 3&4). ‘Fear’ was operationally defined as measurable behavioral freezing (cessation of movement) responses, which were scored manually by an individual blind to the treatment conditions.

### Anatomical tracing

For monosynaptic rabies tracing^50^, CAV2-cre and rgAAV2-mCre (mixed in 1:1 ratio by volume) were injected into the LA/B (AP: -3.0mm, ML: ±5.3mm, DV: -8.6mm, 0.4 uL total injection volume). Rabies helper viruses containing cre-dependent avian TVA receptor AAV9/2-CBA-Flex-TVA950 and rabies glycoprotein AAV9/2-CAGGS-FLEX-H2B-HA-P2A-N2c(G) (mixed in 1:1 ratio by volume) were injected bilaterally into the CnF (AP: -8.4mm, ML:±3.85mm, DV: -5.3mm with 20° angle, 0.3μL total injection volume). Two weeks after the first surgery, EnvA-pseudotyped, glycoprotein gene-deleted rabies EnvA-CVS-N2cΔG-GFP was injected into the CnF (same coordinates as helper AAVs, 0.3μL total injection volume). Rats were sacrificed 10 days after the rabies injection. The brain regions weredefined by manually matching the fluorescent images to the rat brain atlas (Paxinos & Watson). Deep learning-based automated cell segmentation^56^ was used to quantify the labeled cells in each selected region. As a threshold for inclusion of individual brain regions as being retrogradely labeled, the brain regions listed in the figure meet the following criteria: 1. There were GFP-positive cells at the regions appear in all animals and 2. The number of cells in a region consisted of at least 5 cells. The percent of input cells in each brain region is calculated as the number of GFP positive cells divided by number of total GFP positive cells in each brain.

### Histology and Immunohistochemistry

Rats were deeply anesthetized by an overdose of 25% chloral hydrate, then perfused transcardially with 100 mL PBS followed by 50 mL 4% PFA in PBS. The brains were post-fixed in 4% PFA overnight and then moved to 30% sucrose in PBS solution for 48 hrs at 4c. The brains were embedded in Tissue-Tek OCT compound (Sakura Finetek) and frozen for cryostat sectioning (50 um sections). Free-floating brain sections were washed in 0.1 M PBS containing 0.3% Triton-X 100 (PBST), followed by 30-min blocking in PBST containing 2.0% donkey serum (PBSTS). Sections were then incubated in the primary antibodies diluted in PBSTS overnight at 4 °C in the dark. After repeated washing with PBS, sections were incubated for an hour at room temperature in secondary antibodies diluted in PBSTS. After rinsing with PBS, sections were mounted onto slides and cover-slipped with Fluoromount Plus (Cosmobio Co., Ltd.). For c-Fos experiments, rabbit c-Fos primary antibody (Synaptic Systems) was diluted as 1:5000. The secondary antibody was donkey anti-rabbit Alexa Fluor 647 (Invitrogen), diluted as 1:1000. For vGlut2 staining, rabbit vGlut2 primary antibody (Cosmobio Co., Ltd.) was diluted as 1:1000. The secondary antibody was donkey anti-rabbit Alexa Fluor 488 (Invitrogen), diluted as 1:1000. The vGlut2 images were imaged with Olympus FV1000 confocal microscope in z-stack (4 um interslice interval, 3 slices) and maximum intensity z-projection is applied.

For monosynaptic rabies experiments, the primary antibodies were mouse anti-GFP (Thermo Fisher Scientific, 1:2000) and rabbit anti-HA (Cell Signaling Technology, 1:5000). The secondary antibodies were donkey anti-mouse Alexa Fluor 488 (Thermo Fisher Scientific, 1:1000) and donkey anti-rabbit Alexa Fluor 594 (Thermo Fisher Scientific, 1:1000).

### eGRASP experiment

For eGrasp experiments, 0.3 uL of AAV9/2-CWB-yellow pre-eGRASP(p32) was bilaterally injected into CnF, and 0.4 uL of a mixture of AAV1/2-U-mCre and AAV9/2-CAG-Flex-myrmScarlet-I-P2A-post-eGRASP (1:1 ratio, Gift from Bong-Kiun Kaang^48^) was bilaterally injected into LA/B (the same coordinates as described above). After 8 weeks incubation, the animals were perfused and the brain slices containing LA/B were collected. The slices were imaged using an Olympus FV3000 confocal microscope. The mScarlet-expressing cells in LA were located by the 20X objective lens, then the dendrites of the cells were imaged in Z-stack with a 60X oil-immersion lens. The synaptic contacts (the yellow fluorescence signal) on the primary and secondary dendrites were manually identified and counted. To analyze the number of axo-spinous or axo-dendritic puncta, dendritic segments with positive yellow pre-eGRASP signal were included, and the number of puncta was counted for 50 um segments of each dendrite.

### Electrophysiology

To express the inhibitory opsin ArchT in amygdala-projecting cells in the CnF, CAV2-cre and rgAAV2-mCre (mixed in 1:1 ratio by volume) were bilaterally injected into LA/B (AP: -3.0mm, ML: ±5.3mm, DV: -8.6mm, 0.4μL total injection volume); and the AAV9-CAG-FLEX-ArchT-tdTomato was bilaterally injected into CnF (AP: -8.4mm, ML:±3.85mm, DV: -5.3mm with 20° angle, 0.3 μL total injection volume). 6 weeks later, electrode arrays were implanted in LA/B (AP: -3mm, ML: ±5.4mm, DV: -7.0mm), and the optic fibers were implanted bilaterally in CnF (AP: -8.4mm, ML:±3.85mm, DV: -4.8mm with 20° angle). For delivering periorbital electrical shocks, rats were implanted with a pair of insulated stainless steel wires (0.003-inch or 0.076-mm) beneath the skin of the left or right eyelid. The shock wires were positioned in the eyelid contralateral to the recording electrodes. After rats had recovered from electrode implantation surgery, daily screening sessions were conducted in which electrode tips were slowly advanced (40-120 µm per day) into the targeted brain area (lateral amygdala). Neurons were tested for eyelid-shock responses using mild single-shock pulses (1 mA for 2 ms). If no shock responsive neurons were found, the electrodes were advanced.

The recording session began if shock-responsive cells were encountered. Electrodes were connected to a headstage (Neuralynx Inc.) containing 36 unity-gain operational amplifiers (HS36). Spiking activity was digitized at 40 kHz, bandpass-filtered from 300 Hz to 3 kHz and acquired through a Neuralynx data acquisition system. One electrode in a tetrode that did not have any isolated cells during the recording session was used as reference. At the end of the experiment, recording sites were verified histologically with electrolytic lesions by applying 15–20 s of 20-mA direct current through one of the electrodes. Offline single-unit spike sorting was performed using SpikeSort 3D (Neuralynx Inc.).

Animals receive 2 days of handling by experimenters ahead of the recording experiment. In each recording session, animals were placed into a sound-isolating chamber and received eyelid shock US (2ms, 2mA at 7Hz for 1 sec) with or without green laser (532 nm, Laserglow Technologies) illumination to inhibit LA/B projecting CnF neurons. Each recording session consisted of three types of trials: (i) shocks delivered with laser illumination in CnF (from 500 ms before shock onset to 500 ms after shock offset), (ii) shocks delivered without laser, and (iii) laser delivered without shocks. The order of trials was randomized (30 trials in total, with average 90 sec inter-trial interval). Electrodes were advanced 40 µm after the sessions and animals were put back in the homecage. Recording sessions were repeated until the estimated position of electrodes reached basal amygdala.

### Electrophysiology data analysis

Spike data were collected by Neuralynx data acquisition system, and manual spike clustering was performed offline to identify recorded neurons using SpikeSort 3D. Distinct clusters corresponding to specific waveform parameters were manually identified in 3-dimensional feature space. Sorted single-units were further verified using the following criteria: 1) The waveform amplitude varied consistently throughout a recording session, 2) the inter-spike interval was larger than 1 ms (the refractory period of neurons) and 3) the mean spike amplitudes is higher than 70 µV.

For all recorded cells, perievent histograms (PETHs) were derived using timestamps of aversive stimuli. The number of spikes was counted within 50ms time bins. For each neuron, we defined *Ci* as cumulative number of spikes in the *i*th bin across N trials. The average spike count in each bin *Si* was calculated as

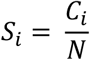

To normalize the neuronal responses, we define the baseline period as the one second period before the shock onset, and calculated the z score as:

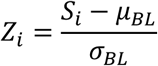

here the mean baseline firing rate is *μBL* and s.d. of baseline firing rate is *σBL*.

The neurons included in the following analysis have firing rates of at least 0.04Hz in the baseline period. Cells were considered shock-responsive if the normalized PSTH reached *Zi* > 2 in at least 1 bin during the one-sec shock period.

### Kinematic analysis

To quantify the animal movement velocity during behavioral experiments, we used DeepLabCut^34^ to track the coordinates of body parts. We built independent network models for behavioral and electrophysiology experiments. The training dataset of behavioral experiments included videos from 6 animals during conditioning, retraining and retrieval sessions. These experiments were done in different contexts and with different patch cord set ups (e.g. conditioning animals had patch cords while in retrieval sessions they did not) and this improved network generalization performance. The training dataset of electrophysiological experiments included videos from 4 recording sessions of each animal. The video frames for training network models were extracted by k-means clustering, 50 frames of each video. We manually labeled the position of the nose, left and right ear, tail stem, and end of tail in each video frame in case the body parts were not clearly visible or partially occluded by the patch cords. 95% of labeled images were used to train the ResNet-50-based neural networks and the other 5% for testing the performance. The network models were refined in three iterations, as the validated performance reached asymptote with limited discrepancy between predicted error in training and test sets (optogenetic experiments: train error: 1.68 pixel, test error: 1.66 pixel with image size: 640x480; electrophysiology experiment: train error: 1.67 pixel, test error: 1.56 pixel with image size: 800x600). Because body parts can be occluded due to wires and patch cords, the coordination of each body part in a video frame was thresholded by the likelihood > 0.95. Kalman filtering and smoothing^82^ was applied to impute data points that fell below the confidence threshold. Escape onset was defined as when velocity during the 1 sec shock shock period reached above 3 standard deviations above average baseline velocity level (1 sec before shock onset, averaged across all trials)

### Mutual information analysis

Animal movements during electrophysiological experiments were recorded with cameras and the velocity motion tracking was described as above. Mutual information (MI)^35^ was calculated to examine the dependence between sensory stimulus states (each individual 2 ms eyelid shock pulse delivered during the 1 sec of US stimulation period, see ‘Electrophysiology’ section) and spiking activity (spike rate) as between whole-body movement (velocity as cm/s) and spiking activity. Mutual information examines conjoint relationships between spiking activity and sensory/motor variables and does not depend on linearity in those relationships. The sensory analysis used each shock pulse and non-shock periods between pulses as discrete, binary states in conjunction with spiking activity for MI analysis. The motor analysis used continuous measures of velocity in combination with spiking activity for MI analysis. The sensory/motor variables were aligned with neuronal responses on a trial-by-trial basis, and the Shannon’s mutual information (*I*) was defined as

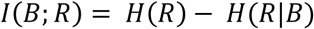

where *H(R)* is the entropy of neuronal responses (*R*); and *H(R|B)* is the conditional entropy of *R* given the behavioral (sensory stimuli or movement speed) variable (*B*), i.e. the random trial-to-trial response variability to the same *B*. To estimate the MI between discrete (for sensory) and continuous (for motor) response measures, we applied Gaussian copula-based analysis to derive the following entropy estimation^83,84^:

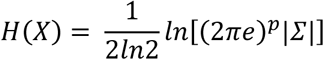

where *p* is the dimensionality of *X* with covariance matrix *Σ*. The upward bias of entropy attributed to limited sampling^59,60^ is corrected as

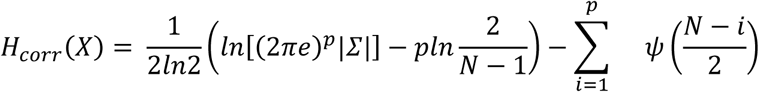

where *N* is the number of trials in a recording session and *Ψ* is the polygamma function (as psi function in MATLAB). The bias-corrected entropy *Hcorr* is used to calculate the Shannon’s MI. To generate control single–unit data with the same number of spikes, the inter-spike interval of the spike trains during the trials was randomly shuffled. Thus, the shuffled spike trains had the same number of spikes and interspike interval distribution as the experimental dataset, but its temporal structure was eliminated.

### Statistics

Statistical analyses were performed using GraphPad Prism 7 (GraphPad Software Inc.) or MATLAB (Mathworks). The sample sizes were not predetermined, but our sample sizes are comparable with previous studies. The behavioral results were analyzed using one-way ANOVA and repeated measures ANOVA with post-hoc correction (Tukey’s or Bonferroni’s multiple comparison tests as determined by GraphPad Prism 7 software). All behavioral data were repeated in multiple groups across days. Anatomical data were collected from several animals with identical rat strain.

## Acknowledgements

We thank Hokto Kazama, Thomas McHugh, Andrea Benucci, Howard Fields and member of the Johansen laboratory for providing valuable feedback on the paper and Bong-Kiun Kang for providing eGRASP plasmids. We acknowledge the support of the RIKEN CBS-OLYMPUS Collaboration Center (BOCC) in our confocal analyses.

## Author contributions

Project conceptualization, LY, JPJ; Experiments & Methodology, LY, JPJ, TO, YK, AU, TD, YI, AJM; Writing – Original Draft, LY, JPJ Writing – Reviewing & Editing, LY, JPJ, YK, TO, AU, AJM, KO; Supervision, JPJ, KO.

---

**Extended Data Fig. 1.**
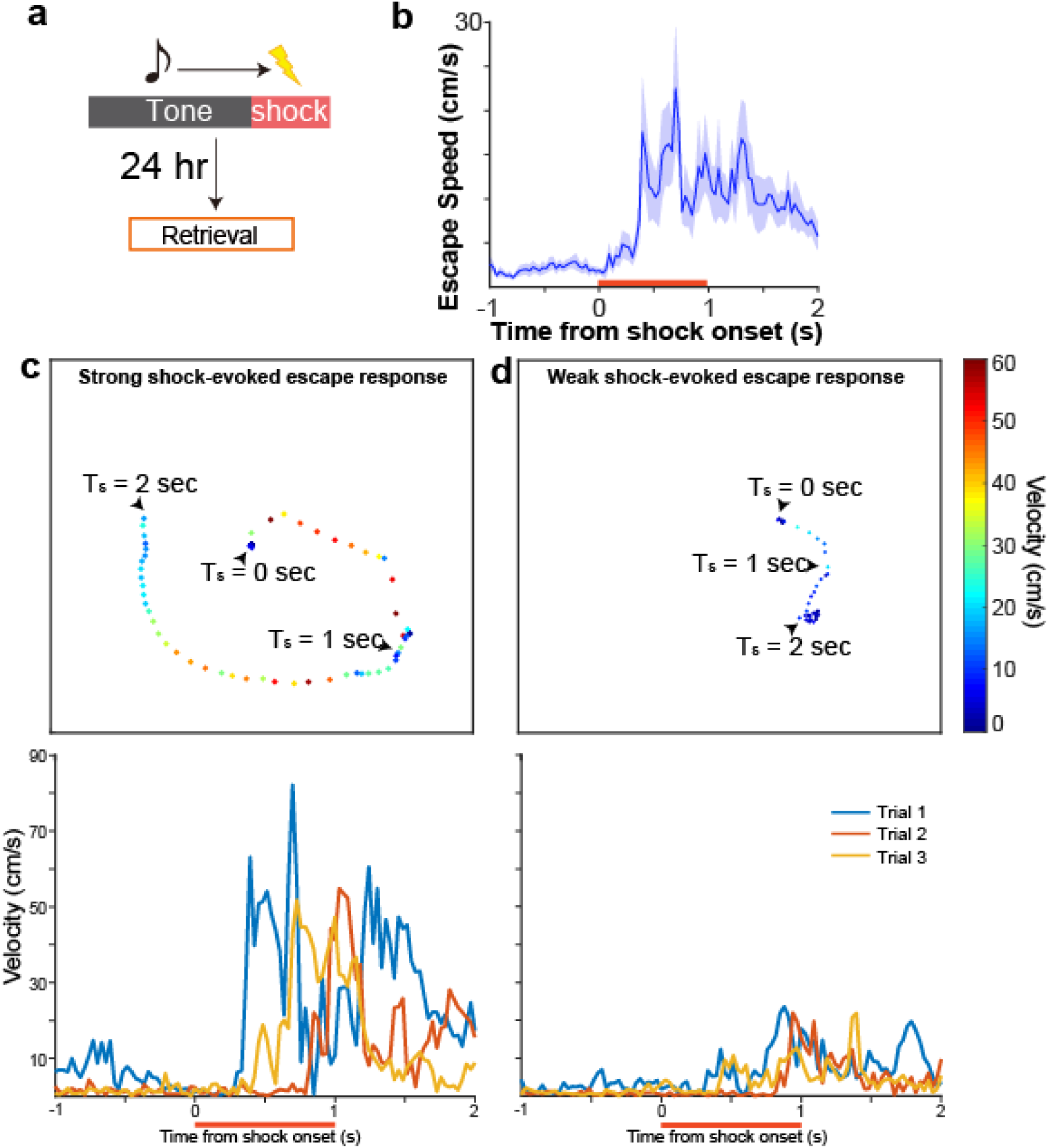
**Characterization of shock-evoked escape behaviors. *a***) fear conditioning paradigm ***b***)Peri-event time histogram (PETH) showing averaged escape velocity (y-axis) before, during and after shock (red bar denotes shock period n=16). Examples of a strong ***c***) and weak ***d***) behavioral escape response to shocks showing speed and distance traveled in space (top graphs) and PETH plotting velocity (y-axis) over time (x-axis) related to the shock onset (bottom graphs). Note considerable variability in the magnitude of escape velocity across trials and animals in lower graphs. Data represent mean ± SEM.

**Extended Data Fig. 2.**
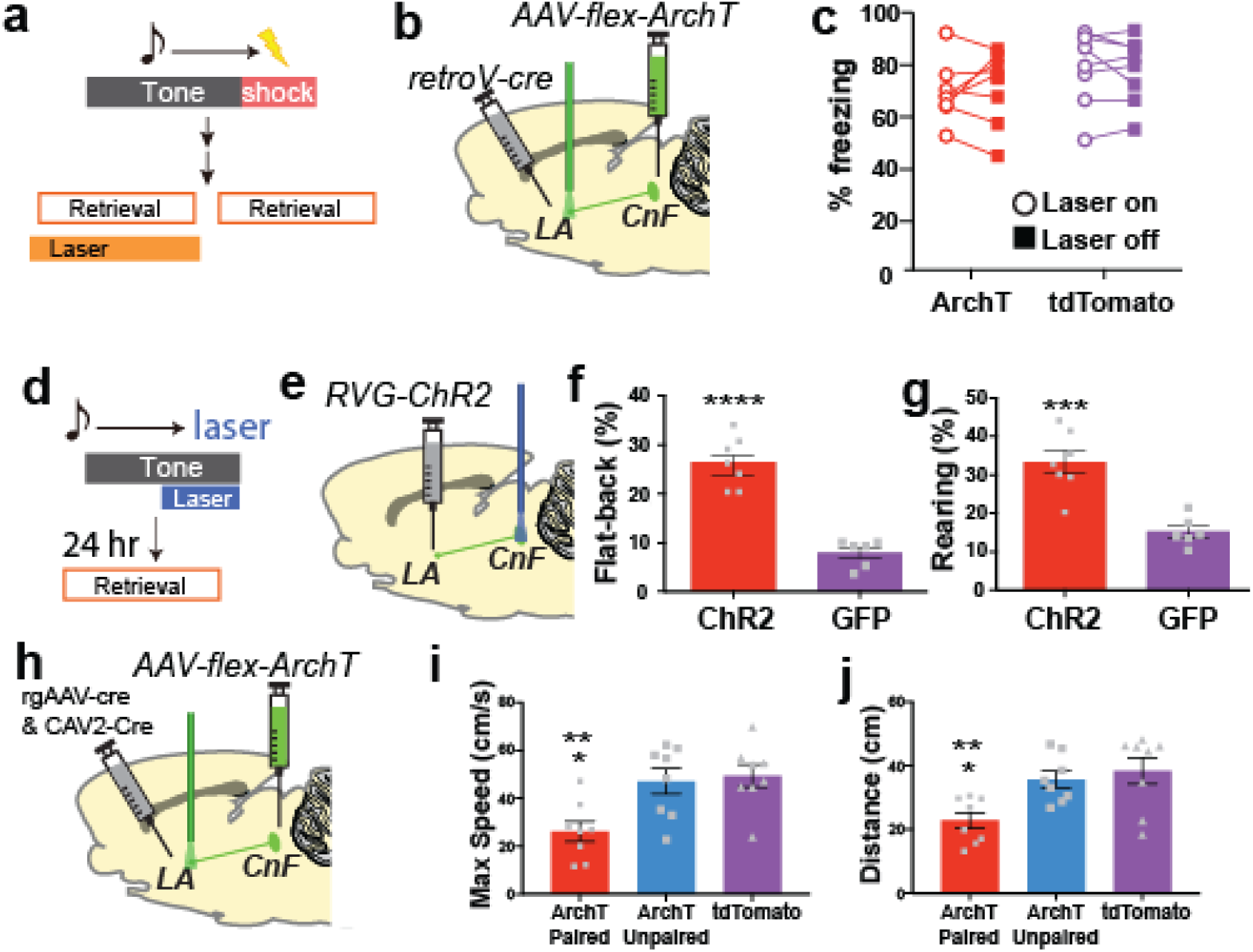
**Effects of optogenetic manipulation of LA/B projecting CnF neurons on various defensive behaviors. *a***) Paradigm for testing effects of CnF-LA/B optogenetic inactivation on the expression of previously learned freezing responses. Laser inhibition of CnF inputs to LA/B during CS period of retrieval occurred in half of CS trials (2 of 4, order was counterbalanced). ***b***) viral/optogenetic strategy. ***c***) Optogenetic inactivation of CnF-LA/B projecting cell bodies has no effect on behavioral freezing (two-way ANOVA, F (1, 42) = 0.07329, *P*=0.7879). Freezing with laser off (closed squares) and on (open circles). ***d***) Paradigm for testing effects of optogenetic stimulation on behavior. ***e***) viral/optogenetic strategy. Optogenetic stimulation produces an increase in flat-back approach (***f***), unpaired t-test, t11=7.781, \*\*\*\**P*<0.0001) and rearing (***g***), unpaired t-test, t11=5.089, \*\*\**P*=0.0003) behaviors. ***h***) viral/optogenetic strategy. ***i-j***) Effects of optogenetic inhibition of CnF terminals in LA on escape response. Inhibition reduced maximum speed (***i***), one-way ANOVA, F(2,21) = 7.228, *P* = 0.0041) and distance traveled (***j***), one-way ANOVA, F(2,21) = 6.762, *P*=0.0054) during shock-evoked escape behaviors.\**P*<0.05, \*\**P*<0.01, \*\*\**P*<0.001, \*\*\*\**P*<0.0001. Data represent mean ± SEM.

